# Adaptive decision making depends on pupil-linked arousal in rats performing tactile discrimination tasks

**DOI:** 10.1101/2022.07.20.500875

**Authors:** Shreya Narasimhan, Brian J. Schriver, Qi Wang

## Abstract

Perceptual decision making is a dynamic cognitive process and is shaped by many factors, including behavioral state, reward contingency, and sensory environment. To understand the extent to which adaptive behavior in decision making is dependent upon pupil-linked arousal, we trained head-fixed rats to perform perceptual decision making tasks and systematically manipulated the probability of Go and No-go stimuli while simultaneously measuring their pupil size in the tasks. Our data demonstrated that the animals adaptively modified their behavior in response to the changes in the sensory environment. The response probability to both Go and No-go stimuli decreased as the probability of the Go stimulus being presented decreased. Analyses within the signal detection theory framework showed that while the animals’ perceptual sensitivity was invariant, their decision criterion increased as the probability of the Go stimulus decreased. Simulation results indicated that the adaptive increase in the decision criterion will increase possible water rewards during the task. Moreover, the adaptive decision making is dependent upon pupil-linked arousal as the increase in the decision criterion was the largest during low pupil-linked arousal periods. Taken together, our results demonstrated that the rats were able to adjust their decision making to maximize rewards in the tasks, and that adaptive behavior in perceptual decision making is dependent upon pupil-linked arousal.

## INTRODUCTION

Adaptive behavior is essential for animals to survive in an ever-varying environment. In perceptual decision making tasks, sensory information is accumulated over time in the central nervous system, eventually leading to a decision to choose one of the alternatives and generating motor commands to indicate the animal’s choice (Smith and Ratcliff, 2004; Gold and Shadlen, 2007; Carandini and Churchland, 2013; Brody and Hanks, 2016; Schriver et al., 2020). The perceptual decision making process is shaped by many factors, including brain state, the gain/loss of each possible decision, task engagement, and external sensory environment (Murphy et al., 2014; Lee and Margolis, 2016; de Gee et al., 2017; Schriver et al., 2018; Cardoso et al., 2019; Kloosterman et al., 2019; Stolyarova et al., 2019; de Gee et al., 2020; Monosov, 2020; Liu et al., 2021; Ashwood et al., 2022). For example, Waiblinger et al. (2019) reported that, in a tactile detection task, rats adjusted their behavioral strategy to maintain a constant payoff in response to changes in probabilistic distribution of whisker deflection amplitudes.

Multiple neuromodulatory systems may exert heavy influences on the cognitive control of adaptive decision making (Doya, 2008). Previous work has suggested that tonic activation of the locus coeruleus - norepinephrine (LC-NE) system plays a critical role in regulating decision making processes with regard to exploring alternatives or exploiting current resources based on uncertainty of available information (Aston-Jones and Cohen, 2005; Yu and Dayan, 2005). On the contrary, phasic activation of the LC-NE system is thought to reset functional networks, and therefore facilitate their reorganization to enable behavioral adaptation in decision making (Bouret and Sara, 2005). It has been postulated that the activity of the cholinergic system is related to expected uncertainty (Yu and Dayan, 2005). The cholinergic system exerts influences on decision making possibly through the widespread ascending projections from the basal forebrain to the cortex (Hasselmo and Sarter, 2011; Pinto et al., 2013; Ballinger et al., 2016; Jepma et al., 2018). Pharmacologically blocking muscarinic receptors suppressed risky decision making in rats performing a gambling task (Betts et al., 2021). Non-luminance-mediated changes in pupil size have been successfully utilized to track rapid fluctuations in cortical state (Reimer et al., 2014; McGinley et al., 2015), and thus are considered as a non-invasive readout of activation of central arousal circuits related to pupil size (i.e. pupil-linked arousal system). Although exact neural circuitry mediating pupil linked arousal remains not fully understood, several lines of evidence suggest that multiple neuromodulatory systems, including the LC-NE and cholinergic systems, contribute to pupil-linked arousal to different extents (Nelson and Mooney, 2016; Reimer et al., 2016; Cazettes et al., 2021). However, how adaptive decision making in response to a varying sensory environment depends on pupil-linked arousal remains unclear.

To address this question, we systematically manipulated the statistics of sensory environment in rats performing a perceptual decision making task. In Go/No-go discrimination tasks, head-fixed rats made perceptual decisions in order to maximize rewards during the tasks. The fraction of trials in which the Go stimulus was presented (i.e. Go stimulus trials) was randomly selected from 0.8, 0.5 or 0.2 for each session. Our data showed that the animals adjusted their behavior in response to the changes in the sensory environment. The response probability to both Go and No-go stimuli decreased as the probability of the Go stimulus being presented decreased. Interestingly, analysis within the signal detection theory framework revealed that the perceptual sensitivity did not vary across the three paradigms. However, the decision criterion increased as the fraction of Go stimulus trials decreased. Simulation results indicated that, to maximize reward per unit time, the decision criterion should increase as the fraction of Go stimulus trials decreases. Therefore, the observed change in decision criterion in the animals indexes their adaptive decision making. Although the animals’ behavior was in line with the optimal trend, the slope of increase in decision criterion across the three paradigms w.r.t. the decrease in fraction of Go stimulus trials was lower than the optimal slope. We compared the slopes of the decision criterions across the three paradigms at high, medium and low pupil-linked arousal levels, and found that the slope was the largest during low pupil-linked arousal periods, suggesting that the animals is able to adjust their decision making processing closer to the optimal level during the low arousal level than the medium and high arousal levels. Drift diffusion modeling further confirmed that the adaptive behavior across the three paradigms was primarily due to adjustment of non-decision time and starting point biases in the decision making process.

## MATERIALS AND METHODS

All experimental procedures were approved by the Columbia University Institutional Animal Care and Use Committee and were conducted in compliance with NIH guidelines. Behavioral studies were conducted using 5 female rats (3 Long Evans and 2 Sprague Dawley, Charles River Laboratories, Wilmington, MA; ∼225-275 g at time of implantation). Animals were single housed after implantation in a dedicated housing facility, which maintained a twelve-hour light and dark cycle.

### Surgical Implantation

The rodent surgery and implantation of head-plates have been described in detail previously (Ollerenshaw et al., 2012; Wang et al., 2012; Ollerenshaw et al., 2014; Schriver et al., 2020). In brief, in aseptic surgeries, anesthesia was induced with a Ketamine/Xylazine cocktail (80/5 mg/kg, IP) or isoflurane (1-3% with a nose cone). Ophthalmic ointment was applied to the eyes throughout the surgery to prevent cornea drying. After the scalp was shaved and hair was removed with depilatory cream, animals were placed in a stereotaxic device using non-penetrating ear bars (David Kopf Instruments, Tujunga, CA). Buprenorphine (Buprenex, 0.03 mg/kg, SC) was administered as an analgesic and Ringers solution (2 mL, SC) was also administered to prevent dehydration. After exposing and cleaning the skull, 8-10 burr holes were drilled in the skull, and stainless steel screws (0-80 thread, McMaster Carr, Robbinsville, NJ) were inserted to anchor a head-plate (Bari et al., 2013; Liu et al., 2021). The wound was then closed with surgical sutures and treated with antibiotic ointment. Antibiotics (Baytril, 5 mg/kg SC) and extra analgesics (Ketoprofen, 5 mg/kg SC) were administered for 5 days postoperatively to minimize the risk of infection. The animals began water restriction and subsequent training following approximately 10 days of recovery from implantation surgery.

### Behavioral Procedures

#### Behavioral apparatus

All behavioral training was conducted in a standard sound and light attenuation chamber (Med Associates, St. Albans, VT). During training, the animals were head-fixed with a custom-made apparatus, in which two pneumatic cylinders on either side of the head that were fixed with ball bearings aligned with grooves in the head plate to hold the animals’ head (Rodenkirch et al., 2019; Liu et al., 2021). A 1 mL syringe body which served to deliver water was mounted to a flexible beam and placed directly in front of the animal. A piezoelectric force sensor was bonded to the flexible beam to measure voltage swings resulting from animals licking the syringe. The sensor’s output was sampled by a DAQ card (PCI-6259, National Instruments, Dallas, TX) at 1 kHz.

Precise tactile stimuli were delivered via a multilayer piezoelectric bending actuator (PL140; Physik Instrumente, Germany) driven by a high-voltage amplifier (OPA452; Texas Instruments, Dallas, TX). Whiskers were placed in a short glass capillary pipette approximately 15 mm long with an outer diameter of 1 mm and an inner diameter of 0.5 mm (AM Systems, Carlsborg, WA). The pipette was bonded to the end of the piezo actuator and placed 8 mm away from the right snout. The whisker that received tactile stimuli was chosen with respect to its thickness between C2, C3 and D2, and the same chosen whisker was used in all behavioral sessions for each animal. The chosen whisker was slightly trimmed to facilitate the insertion into the pipette.

To mask possible auditory cues, a buzzer (bandwidth: 16 Hz – 10 kHz) delivering white noise-masking sound was placed next to the whisker stimulator. Onset tone (6 kHz), reward tone (3 kHz), and timeout tones (16.5 kHz) were delivered by a speaker installed in the chamber. Animals were remotely monitored with a CCD camera, and an infrared LED was placed in the chamber for illumination during the task. Control of the behavioral task and sampling of animals’ behavioral responses were performed by custom-programmed software running on a MATLAB xPC target real-time system (Mathworks, Natick, MA). All behavioral data was sampled at 1 kHz and logged for offline analyses.

#### Tactile stimulus

Whisker stimuli used were sinusoidal waveforms of 8 Hz and 4 Hz (0.5 s, 1 mm amplitude), with the 8 Hz stimulus randomly assigned as the Go stimulus and the 4 Hz stimulus as the No-go stimulus. The probability of the Go stimulus being presented was designated as either 80%, 50%, or 20% for each session.

#### Pupillometry recording

Recording of the pupil contralateral to the whisker deflection was made using a custom-made pupillometry system (Liu et al., 2017), which were triggered at 20 Hz by the xPC target real-time system (Mathworks, MA) that controlled the behavioral task. Pupil images were streamed to a high-speed solid-state drive for offline analysis. To extract pupil size, the pupil contour was segmented using the DeepLabCut toolbox (Mathis et al., 2018). A training set consisting of two hundred frames recorded across different sessions was selected. Each frame had 12 evenly distributed points labeled surrounding the pupil manually, and the images were cropped to enable a higher training accuracy. The ResNet50 deep network was used to analyze the video clips from all sessions after the training. The automatically labelled points were fit with circular regression and the pupil size was computed as the area bounded by the contour. 5% of all images were randomly selected for inspection to validate the accuracy of the software. Pupil sizes during blinks were interpolated with values before and after blinks (Nassar et al., 2012; Liu et al., 2017). The pupil size was low-pass filtered with a fourth-order non-causal filter with a cutoff frequency of 3.5 Hz.

#### Training and the Go/No-go Discrimination Task

Water deprivation schedule and procedures of head-fixation habituation were the same as our previous work (Schriver et al., 2018; Schriver et al., 2020; Liu et al., 2021). Briefly, restriction of access to water was used to motivate animals during the tasks. However, during the behavioral task, correct responses to a Go stimulus were rewarded with ∼60 uL aliquots of water. Because the number of possible rewarding trials (i.e. Go stimulus trials) was different across the three paradigms, supplemental water was given before returning the animals to the animal facility to ensure their daily water intakes were the same across all training days. The weight of the animals was measured and logged immediately after the task.

The onset of each trial was indicated by a brief “trial onset tone” (300 ms, 6 kHz), followed by a random delay (1 to 3.5 s uniform distribution) (Figure 1B). To discourage the animal from impulsively licking, the last 1 s of the waiting period was a designated “no lick” period, during which any premature licks would result in an additional delay in stimulus presentation pulled from a 1 - 2.5 s uniform distribution (Stuttgen and Schwarz, 2008; Ollerenshaw et al., 2014). The stimulus for each trial could be either a Go stimulus or a No-go stimulus, but the fraction of Go stimulus trials was randomly selected from 0.8, 0.5 and 0.2 for each session, resulting in three behavioral paradigms. Licking within a window of opportunity (1.3 s) following a Go stimulus resulted in a brief “reward tone” (300 ms, 3 kHz) accompanied by a water reward, whereas licking within the window of opportunity following a No-go stimulus triggered a “timeout tone” (5 s, 16.5 kHz) which began a 10 s timeout period. CR and miss behavioral outcomes were neither rewarded nor penalized. A 6 s inter-trial period followed the end of the window of opportunity for CR and miss trials, water reward for hit trials, and timeout period for false alarm (FA) trials. Across all 5 animals, 260 sessions were performed and 70516 trials were recorded. Pupillometry was recorded in 165 sessions.

**Figure 1.**
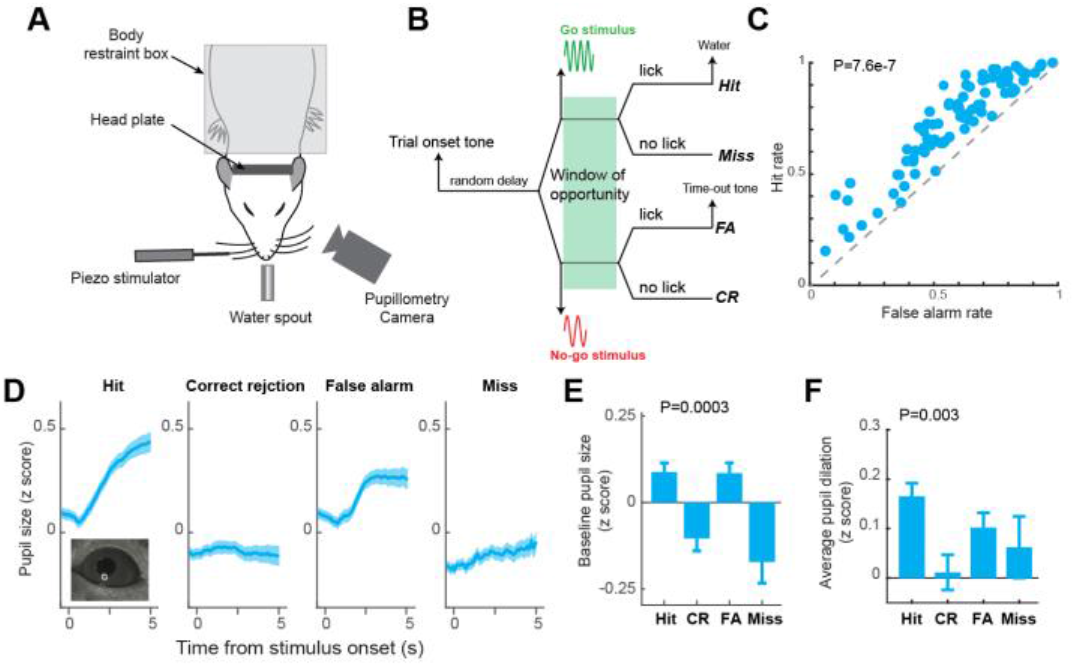
Behavioral performance and pupil dynamics during a tactile decision making task. **A)** Experimental setup. **B)** The diagram of a Go/No-go perceptual decision-making task. **C)** Hit rate was significantly higher than false alarm rate across sessions for the animals. **D)** Pupil dynamics around stimulus presentation associated with the four behavioral outcomes. **E)** Baseline pupil size associated with the four behavioral outcomes. **F)** Pupil dilation associated with the four behavioral outcomes.

### Data Analysis

All data analyses were first conducted on individual sessions. Grand averages and standard errors of means were then calculated across sessions for analysis and presentation. For each session, the first 20 trials were excluded due to the time required to adjust the pupillometry camera.

#### Behavioral Performance

Response probabilities for each session were calculated as the hit rate (HR, i.e. number of hit trials/number of S+ trials) and FA rate (FAR, i.e. number of FA trials/number of S-trials). These were used to calculate perceptual sensitivity (d’) and decision criterion as

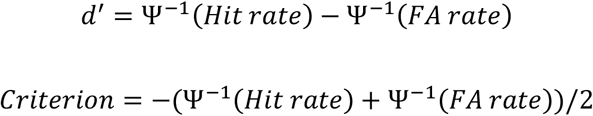

Where Ψ^−1^ is inverse of the *cumulative Gaussian* distribution.

For analyzing response probabilities, perceptual sensitivity, and decision criterion versus percent of maximum baseline, each session’s baseline range was first computed and then evenly broken into 20 bins, each trial was sorted into one of the bins, and HR, FAR, d’, and criterion were calculated for each bin. The loglinear approach was utilized to allow for calculating d’ and criterion in bins where HR or FAR equaled 1 or 0, where 0.5 was added to the number of hits and FAs and 1 was added to the number of S+ and S-presentations prior to calculating HR and FAR (Stanislaw and Todorov, 1999).

Reaction times were computed as the time from stimulus onset, which is when the window of opportunity began, until the first lick response within the window of opportunity. Reaction times were only computed when a response was logged within the window of opportunity, i.e. for hit and FA trials, but not miss or CR trials.

#### Pupil dynamics

Pupil sizes were first Z-scored for each session prior to further analyses. Pupil sizes were aligned by stimulus onset. Stereotypical pupil responses for each behavioral outcome were calculated as the average pupil size at each time point 0.5 s preceding stimulus onset to 5 s following stimulus onset. Average baselines and dilations were calculated from these averaged stereotypical responses for each behavioral outcome for each session. Baseline pupil sizes were computed as the average of the pupil sizes in the 0.5 s preceding the stimulus, while dilations were calculated as the mean value of the pupil sizes from stimulus onset to 5 seconds post stimulus onset, minus the pupil baseline. To calculate the percent of maximum pupil baseline, all baselines were normalized for each session.

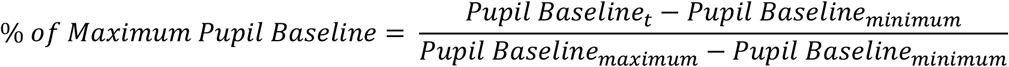

#### Simulation to determine optimal decision criterion

To determine optimal decision criterion, we simulated the behavior of rats with different decision criterion in the three paradigms based on the signal detection theory and computed water reward per unit time for each decision criterion in each paradigm. For each simulated session, the probability of S+ trials was set at either 20%, 50%, or 80%. For a given decision criterion, on an S+ trial, a random number was drawn from a normal distribution with mean = 0.52, which is the mean perceptual sensitivity across the three paradigms, and variance = 1. If the random number was greater than the decision criterion, a hit was logged. Otherwise, a miss was logged. For a No-go trial, a random number was drawn for a normal distribution with mean of 0 and variance of 1. Either a false alarm or correct rejection was logged, depending upon if the random number was greater than the decision criterion. The duration of a hit trial was composed of a random waiting period (from a 1 - 3.5 s uniform distribution), a mean response time, and a 6 s inter-trial interval, while the duration of a miss or correction rejection trial is composed of a random waiting period (from a 1 - 3.5 s uniform distribution), a 1.3 s of window of opportunity and a 6 s inter-trial interval. The duration of a false alarm trial is the sum of a random waiting period (from a 1 - 3.5 s uniform distribution), a mean response time, a 10 s timeout, and a 6 s inter-trial interval. For each paradigm, we simulated 15,000 trials for each decision criterion, and the decision criterion leading to the maximal water reward per unit time was considered as the optimal decision criterion. Note that water rewards resulted only from hit responses. We repeated the simulation 20 times to estimate the variance of the optimal decision criterion for each paradigm.

#### HDDM modeling

Drift diffusion models (DDM) were used to quantify possible differences in the parameters of decision making across the thee behavioral paradigms (Wiecki et al., 2013). We compared four DDM models with various parameter constraints (Delis et al., 2018; Schriver et al., 2020; Delis et al., 2022). The first model assumed an unbiased starting point, i.e. the start point is the middle of decision boundary 0.5a or z = 0.5 (Figure 6A). For both Go and No-go conditions, the absolute value of the drift rate is the same. The second model assumed that the starting point can be biased, but that the absolute value of the drift rate is the same for both Go and No-go conditions. The third model assumed an unbiased starting point just like the first model, but the drift rate was allowed to be different between the Go and No-go conditions. The fourth model assumed biased starting points and different drift rates for the Go and No-go conditions.

To minimize the risk of overfitting, we calculated the deviance information criterion (DIC) value for each model. Because DIC measure is a tradeoff between goodness of fit and number of free parameters for Bayesian models, we only considered the model with the least DIC as the optimal fit. Each parameter of the model had three group-level distributions corresponding to the three behavioral paradigms and individual-level distributions for each animal.

## RESULTS

To test how animals adaptively change their behavior in perceptual decision making tasks in response to changes in sensory environment, we trained head-fixed rats to perform tactile decision making tasks using a Go/No-go discrimination paradigm (**Figure 1A**) (Schriver et al., 2018; Liu et al., 2021). In these tasks, the rats were required to make decisions to respond or withhold response after a tactile stimulus was presented (**Figure 1B**). In the initial training sessions, Go stimulus (S+ stimulus, 0.5 s 8 Hz whisker stimulation), which the animal is trained to respond to for rewards, was randomly presented in 50% of trials, while No-go stimulus (S-stimulus, 0.5 s 4 Hz whisker stimulation), to which the animal is trained to withhold response to avoid time-out, was presented in the rest of the trials. Animals had significantly higher response probability to S+ than to S-stimulus in these sessions (0.72±0.019 vs 0.58±0.02, p<7.6e-07, paired t-test, mean ± SEM unless otherwise noted, **Figure 1C**), indicating that the animal understands the requirement. Moreover, consistent with our previous work, the pupil size of the rats fluctuated throughout the sessions. The pupil dynamics around stimulus presentation were different across the four possible behavioral outcomes (i.e., hit, correct rejection, false alarm, miss) (**Figure 1 D&E**). Baseline pupil size before stimulus onset was higher for hit and false alarm trials than for correct rejection and miss trials (hit: 0.08±0.023; correct rejection: −0.085±0.036, false alarm: 0.074±0.027, miss: −0.14±0.07, p=0.0003, ANOVA test). Task evoked pupil dilation was largest for hit trials, followed by false alarm trials, miss trials, and correct rejection trials (hit: 0.156±0.015; false alarm: 0.096±0.018; miss: 0.07±0.034; correct rejection: 0.0095±0.022; p=0.003; ANOVA test).

To test whether the animals adaptively changed their behavior in response to changes in sensory environment, we systematically manipulated the statistics of sensory signals. In our experiments, we used three paradigms in which fractions of S+ trials, i.e. trials on which S+ stimulus was presented, were set at 20%, 50%, and 80%. Each session was randomly assigned with one paradigm and its corresponding fraction of S+ trials. We found that animals adaptively changed their response rate in response to changes in the fraction of S+ trials for each session (**Figure 2A**). In general, both hit rate and false alarm rate decreased as the fraction of S+ trials decreased from 80% to 20% (0.8574±0.0021 vs 0.723±0.002 vs 0.552±0.003, p<9e-16, ANOVA test; false alarm rate: 0.7622±0.0025 vs 0.578±0.002 vs 0.398±0.003, p<1.9e-22 ANOVA test) (**Figure 2B**). Using the signal detection theory framework (Wang et al., 2010; Ollerenshaw et al., 2012; Bari et al., 2013; Zheng et al., 2015), we found that the perceptual sensitivity of the animals was not affected by the changes in fraction of S+ trials, as there was no significant difference across the three paradigms with different fractions of S+ trials (0.536±0.042 vs 0.510±0.029 vs 0.518±0.045, p=0.88, ANOVA test) (**Figure 2C**). However, we found that the animals systematically adjusted their decision criterion from more liberal (i.e. more negative decision criterion) to more conservative (i.e. less negative decision criterion) in responses to the decrease in the fraction of S+ trials (−1.139±0.08 vs −0.483±0.064 vs 0.045±0.085, p<3.1e-20, ANOVA test) (**Figure 2D**).

**Figure 2.**
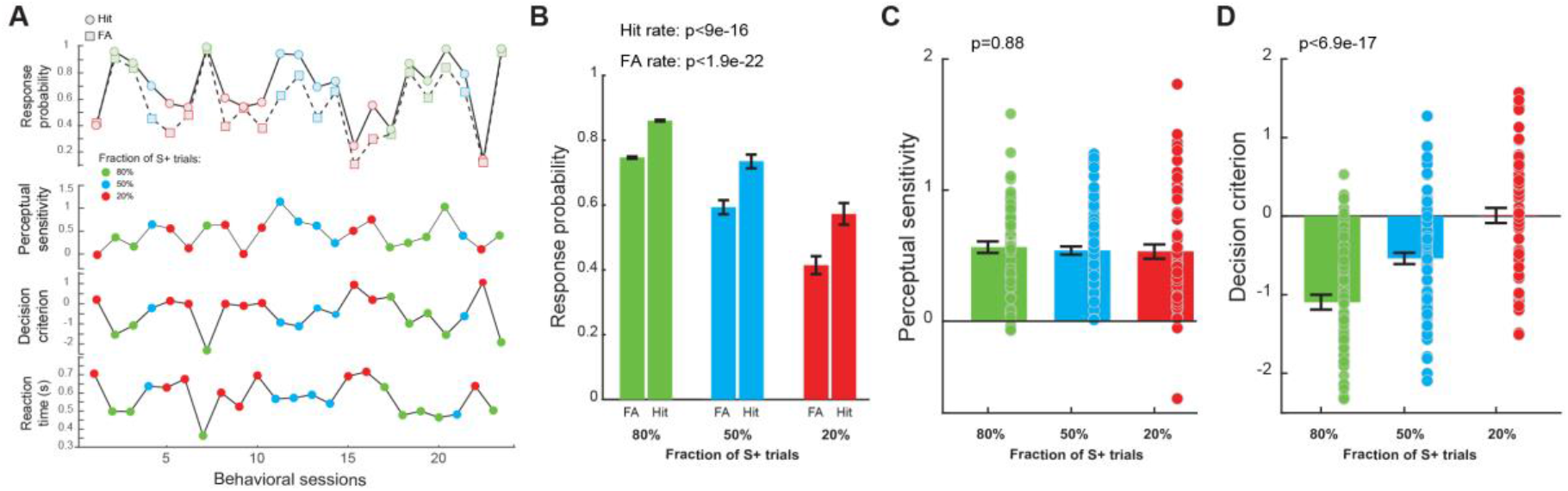
Adaptive behavior in response to change in the statistics of sensory environment. **A)** Example adaptive behavior in response to change in the probablity of S+ trials. **B)** Response probability across the three paradigms. **C)** Perceptual sensitivity remained the same across the three paradigms. **D)** Decision criterion significantly increased as the probability of S+ trials decreased.

Because animals only received water rewards on hit trials during the task, animals could accumulate less water intake in sessions with the fraction of S+ trials being 0.2, compared to the other two paradigms. Although we gave each animal supplemental water right before we returned them to the animal facility on each training day to ensure they have the same daily water intake across sessions with different paradigms, it is still possible that the difference in level of thirst during the task may have resulted in the changes in the response probability to both Go and No-go stimuli. However, if the animal is thirstier, it is plausible that it should have high response rates in an attempt to get more water rewards. However, this is in contrary to what we experimentally observed (**Figure 2B**). To further rule out this possibility, we calculated the percent of trials in which the animal impulsively licked (i.e. licked between trial onset tone and stimulus presentation). We reasoned that if animals were thirstier in sessions where 20% of the trials were S+ trials, they would tend to lick more impulsively, leading to a higher impulsive licking rate. However, our data suggested this was not the case. The fraction of impulsive licking trials was significantly smaller for sessions where 20% of the trials were S+ trials, compared to the other two paradigms (0.174±0.019 vs 0.134±0.0106 vs 0.062±0.0055, p<1.06e-8, ANOVA test) (**Figure 3A**), suggesting that the changes in response probability resulted from cognitive processing in responses to changes in probability of S+ trials, rather than the level of thirst. To further support this notion, we found that the reaction time monotonically increased with the decrease in fraction of S+ trials (0.4838±0.012 s vs 0.594±0.01 s vs 0.702±0.011 s, p<1.82e-29, ANOVA test) (**Figure 3B**). Furthermore, decision criterion was positively correlated with reaction time (p<4.23e-25) (**Figure 3C**). As we expected, our data showed that decision criterion was negatively correlated with percent of impulsive licking trials (p<2.23e-15) (**Figure 3D**). Taken together, these results suggest that the adaptive behavior of the animals that we observed in our experiments was due to higher-level cognitive processing of the statistics of sensory environment, rather than low-level physiological needs such as thirst.

**Figure 3.**
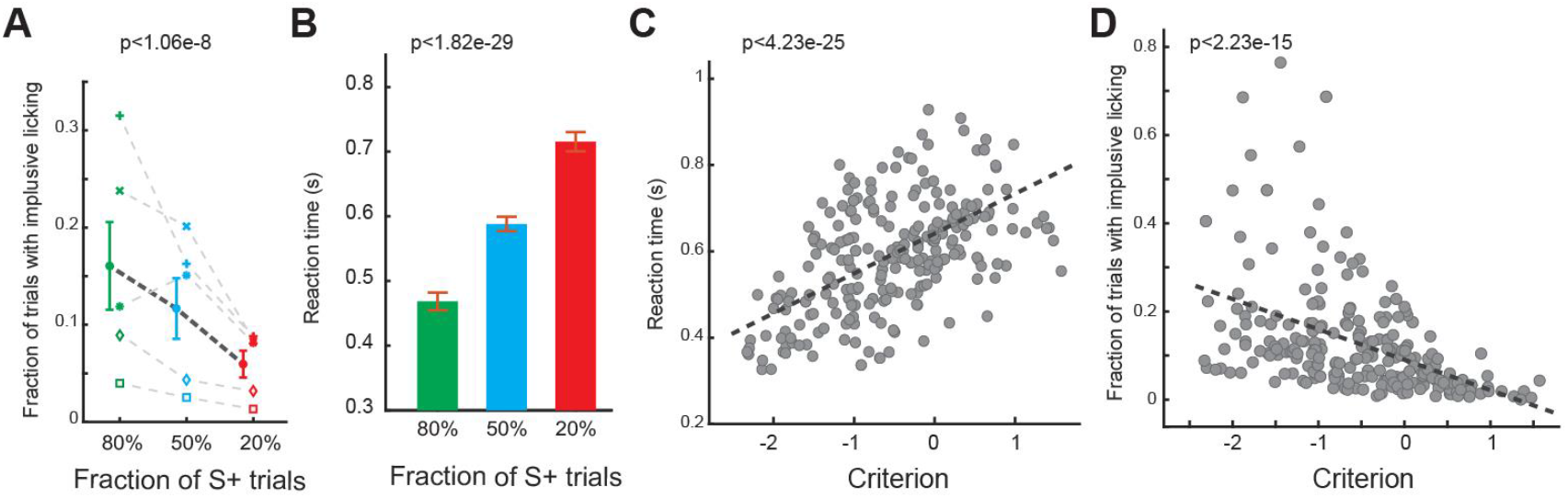
Changes in impulsive licking rate and reaction time in the three paradigms. **A)** Impulsive licking rate decreased along with the decrease in probability of S+ trials. **B)** Reaction time significantly increased as the probability of S+ trials decreased. **C)** Reaction lime is positively correlated with decision criterion. **D)** Impulsive licking rate is negatively correlated with decision criterion.

We have previously showed that perceptual decision making depended on pupil-linked arousal (Schriver et al., 2020). We then examine if pupil dynamics were different during adaptive decision making across the three paradigms. Indeed, the pupil dynamics around stimulus presentation were significantly different between the three paradigms for the four behavioral outcomes (**Figure 4A**). We found there was a dramatic difference in task-evoked pupil dilation between the three paradigms for hit trials (0.0554±0.02 vs 0.156±0.015 vs 0.36±0.03, p<1.9e-15, ANOVA test), and to a lesser degree for false alarm trials (0.098±0.022 vs 0.096±0.018 vs 0.173±0.0195, p<0.01, ANOVA test), but not for the other two behavioral outcomes (correct rejection: 0.0525±0.037 vs 0.009±0.028 vs 0.154±0.014, p=0.45, ANOVA test; miss: 0.0595±0.0466 vs 0.07±0.035 vs 0.04±0.043, p=0.90, ANOVA test) (**Figure 4B**).There is a significant change in baseline pupil size between the three paradigms for false alarm trials (0.0142±0.029 vs 0.074±0.07 vs 0.15±0.04, p<0.017, ANOVA test), while there was not a significant difference in baseline pupil size between the three paradigms for the other three behavioral outcomes (hit: 0.049±0.017 vs 0.08±0.023 vs 0.113±0.04, p=0.32, ANOVA test; correct rejection: −0.11±0.062 vs −0.085±0.036 vs −0.183±0.034, p=0.27, ANOVA test; miss: −0.0194±0.087 vs −0.1426±0.007 vs −0.183±0.072, p=0.32, ANOVA test) (**Figure 4C**). Interestingly, in paradigms with the fraction of S+ trials being 0.8 and 0.5, there is a profound inverted-U or U shaped relationship between baseline pupil size and hit/false alarm rates, decision criterion, and perceptual sensitivity (**Figure 4D&E**). However, this inverted-U or U shaped relationship between baseline pupil size and hit/false alarm rates, decision criterion, and perceptual sensitivity is less conspicuous for the paradigm in which the fraction of S+ trials is 0.2 (**Figure 4F**).

**Figure 4.**
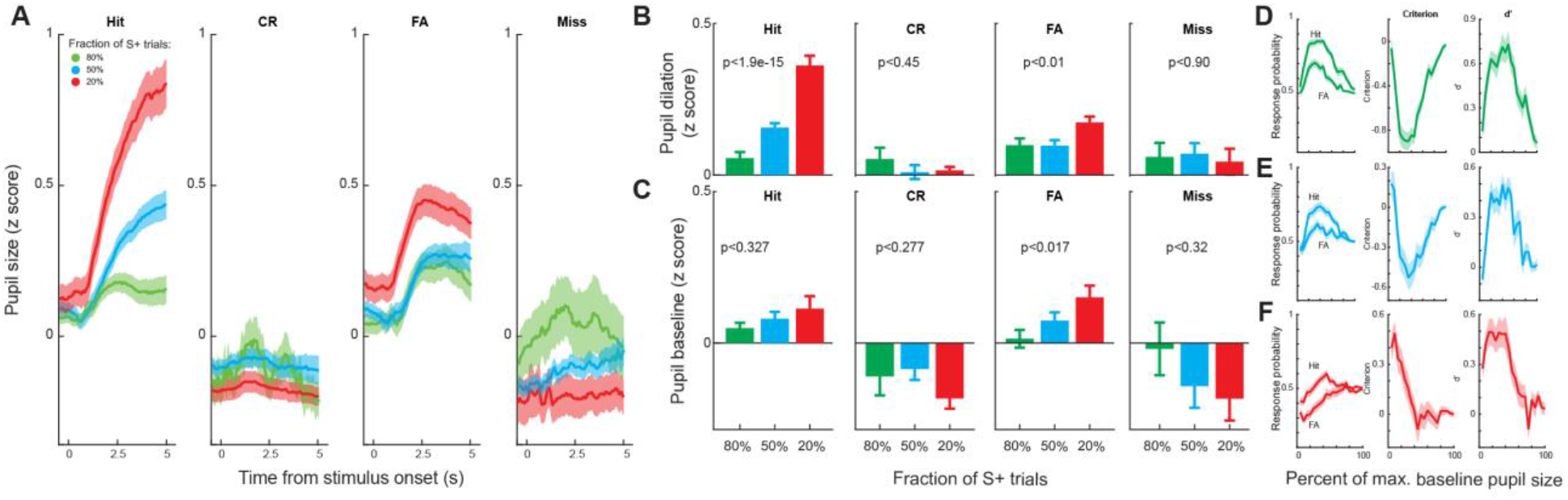
Pupil dynamics in the three paradigms. **A)** Pupil dynamics around stimulus presentation associated with the four behavioral outcomes in the three paradigms. **B)** Baseline pupil size associated with the four behavioral outcomes in the three paradigms. **C)** Pupil size associated with the four behavioral outcomes in the three paradigms. **D-F)** The relationship between response probability/decision criterion/perceptual sensitivity and baseline pupil size in the three paradigms.

How were the pupil dynamics related to the adaptive behavior? Our data demonstrated that task-evoked pupil dilations were different across the three paradigms, and that the animals mostly adjusted their decision criterion while maintaining the same perceptual sensitivity across the three paradigms. To determine the optimal decision criterion for each paradigm, we simulated the water reward per unit time with different decision criteria for each paradigm using our experimental parameters. If a decision criterion is too negative, the animal will be liberal in making Go decisions. Consequently, they will encounter many false alarms, and thus a substantial portion of the task will be in the time-out period. On the contrary, if the animal is too conservative in making Go decisions and sets the decision criterion to be a large positive value, the animal will falsely reject many S+ stimuli, resulting in a low hit rate and less water intake throughout the task period. Our simulation results indicate that the optimal decision criterion was significantly smaller than the ones that we observed experimentally. For the paradigm with fraction of S+ trials being 0.8, the optimal decision criterion and observed decision criterion were −3.215±0.107 vs −1.137±0.084 (p<0.2.35e-20, t-test). Similarly, the optimal and observed decision criterion were −1.535±0.033 vs −0.479±0.065 (p<1.43e-10, t-test) and −0.93±0.0275 vs 0.0636±0.086 (p<6.5e-08, t-test) for the two other paradigms, respectively (**Figure 5A**). We further examined if the decision criterion is dependent upon pupil size within each paradigm. We found that in the paradigm where 80% of trials were S+ trials, the decision criterion increased monotonically with baseline pupil size (p<0.0025). However, this trend did not hold for the other two paradigms (**Figure 5B**).

**Figure 5.**
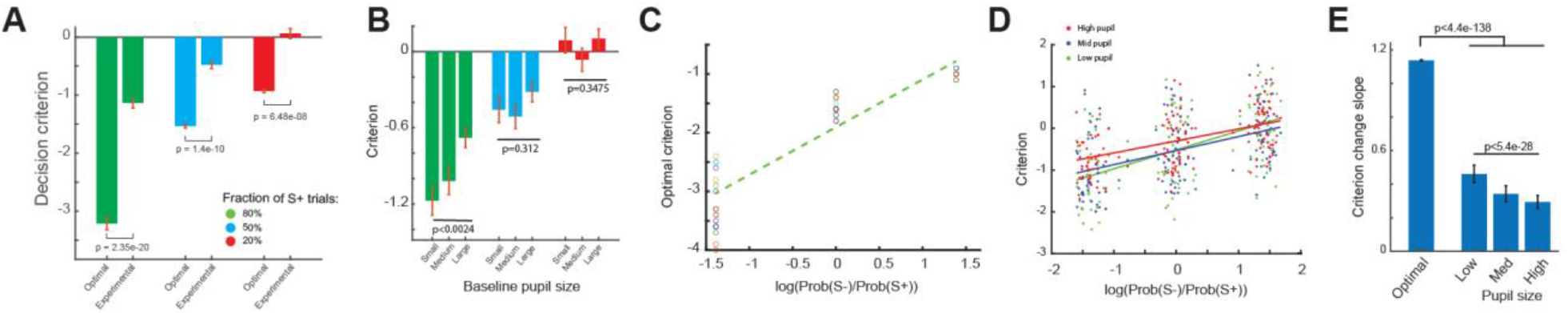
Optimal adjustment of decision criterion is dependent upon pupil size. **A)** Optimal and experimentally observed decision criterion in the three paradigms. **B)** Decision criterion of the animals at different pupil-linked arousal levels. **C)** Optimal adaptation of decision criterion in response to changes in probability of S+ trials. **D)** Adaptation of decision criterion in response to changes in probability of S+ trials in the animals at different pupil-linked arousal levels. **E)** Optimal and experimentally observed adaptation of decision criterion in response to changes in probablity of S+ trials.

Although these results indicated that the animals were sub-optimal in terms of their adaptive behavior, the adaptive change in decision criterion observed in our experiments was in line with the optimal adjustment of decision criteria. We therefore compared changes in decision criteria across the three paradigms, measured as the slope of decision criterion across the three paradigms, between the optimal decision making case (i.e. simulation) and the real case (i.e. experiments). We found that the change in optimal decision criteria in response to changes in paradigms, i.e. slope of optimal decision criterion across the three paradigms, was much stiffer than the experimentally observed slope (1.14 vs 0.355) (**Figure 5C**). We further examined if the adjustment of decision criterion in response to changes in sensory environment depended on pupil-linked arousal indexed by pupil size. To this end, we grouped trials of each session into three groups based on the baseline pupil size, and calculated decision criteria for trials within each group (**Figure 5D**). We found that there was a systematic change in the slope along the baseline pupil size, with the slope being largest during small baseline pupil size (p<5.4e-28, ANOVA test). However, the slopes for all pupil size dependent groups were significantly smaller than the slope of optimal decision criteria (p<4.4e-138, ANOVA test) (**Figure 5E**).

We further used the drift diffusion model (DDM) to quantify the extent to which the other parameters of decision making, including non-decision time, decision boundary, drift rate and initial bias, were affected by the changes in sensory environment (**Figure 6A**). To this end, we used a Bayesian approach to estimate the distributions of decision making parameters at the group level for each paradigm (Wiecki et al., 2013), and P_p|D_ is used to refer to the proportion of posteriors from Bayesian inference, supporting the working hypothesis that there is a difference between the paradigms at the group level. We first calculated the Deviance Information Criterion (DIC) value of the four variants of the hierarchical DDM for our behavioral data (see Methods). Since DIC balances between a goodness-of-fit of the model and additional free model parameters, we used the model with the lowest DIC value (**Figure 6B**). This model generated a similar distribution of reaction times to those measured experimentally (**Figure 6C**). HDDM results suggested a significant difference in non-decision time and initial bias among the three paradigms (P_p|D_≈1) (**Figure 6D&E**). However, for the decision boundary, there is only a significant difference between the paradigm with 80% S+ trials and both paradigms with 50% and 20% S+ trials (P_p|D_≈1), and there is no difference between the paradigm with 50% S+ trials and the paradigm with 20% S+ trials (P_p|D_=0.2) (**Figure 6F**). We failed to find significant differences in drift rate across the paradigms (P_p|D_>0.05) (**Figure 6G**).

**Figure 6.**
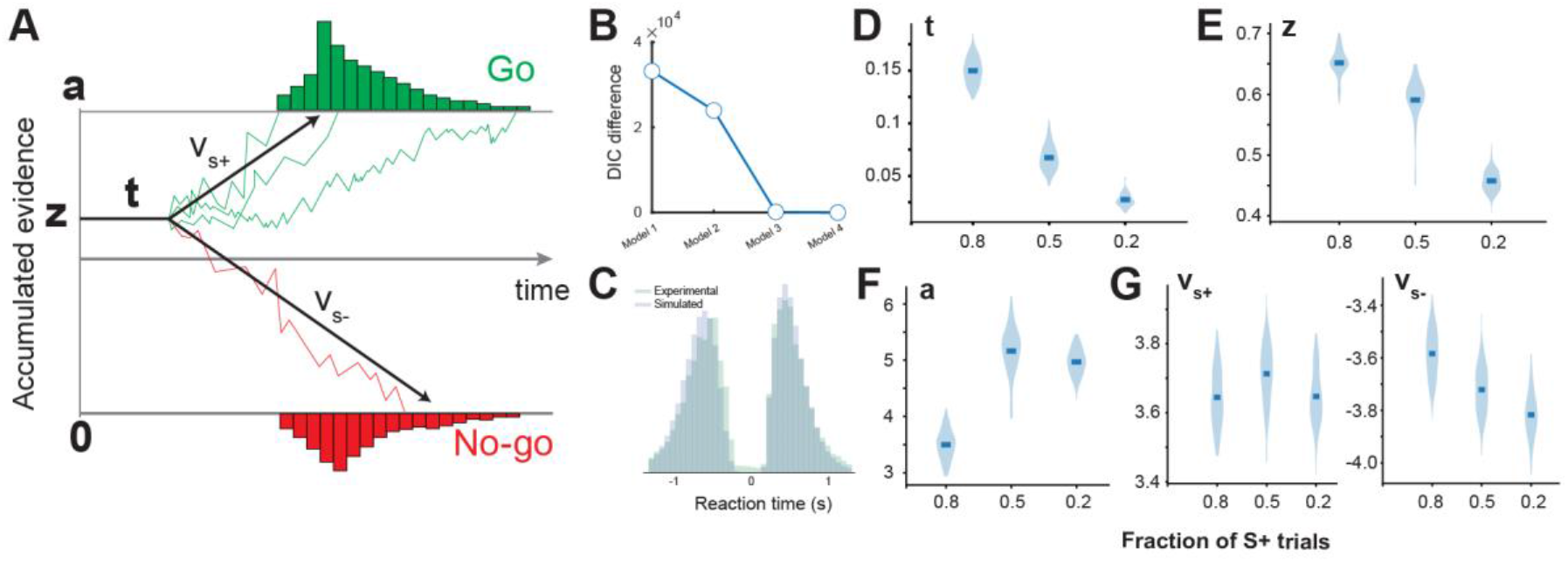
HDDM modeling of perceptual decision making in the three paradigms. **A)** Diagram of the drift diffusion model. **B)** DIC difference between the four alternative models. **C)** Reaction time distributions were captured by the HDDM. **D-G)** Violin plots showing the posterior estimates of the HDDM parameters for the three paradigms.

## DISCUSSION

Our previous work investigated how pupil-linked arousal modulates behavioral performance (Schriver et al., 2018) and the extent to which arousal systems indexed by electrocardiograph signals and pupil size differently modulate perceptual behavior (Liu et al., 2021). The present study was designed to allow us to investigate adaptive behavior in rats performing perceptual decision making tasks. Previous work has demonstrated adaptive behavior in response to changes in rewards and stimulus properties (Cardoso et al., 2019; Waiblinger et al., 2019). In this study, we manipulated the probability of Go and No-go stimuli and characterized the animals’ adaptive decision making at different pupil-linked arousal levels. The results present several novel findings. First, we showed that the animals became more liberal in making a Go decision when the probability of S+ stimulus increased. This adaptation is in line with the optimal adaptation to maximize rewards during the task (**Figure 2&5**). Second, task evoked phasic pupil-linked arousal is higher when probability of S+ is low (**Figure 4**). Finally, adaptive decision making is dependent upon pupil-linked arousal (**Figure 5D&E**). Consistent with human results (Kahneman and Tversky, 2000), our animals performed sub-optimally in adjusting their decision criterion in response to varying probabilities of S+ trials. For all three paradigms, the animals could have received more water rewards if they were more liberal in making Go decisions in the behavioral tasks (**Figure 5A**). One possible explanation could be that the tone indicating the onset of timeout period became aversive to the animals, increasing the cost of false alarms.

We systematically manipulated the probability of S+ trials, varying from either 20%, 50% or 80% in the experiments, as a means to probe adaptive decision making. Uncertainty of sensory inputs and rewards has been shown to impose effects on neural computation and decision making (Yu and Dayan, 2005; Orbán and Wolpert, 2011; Kepecs, 2013; Payzan-LeNestour et al., 2013; Urai et al., 2017; Stolyarova et al., 2019). It is important to note that the uncertainty of sensory stimuli is theoretically the same for the two paradigms in which the probability of S+ trial is 80% and 20% in our experiments. If the uncertainty plays a critical role in adaptive decision making and evoking pupil dilation in our experiments, we would expect to see the same adaptive behavior and pupil dynamics in those two paradigms. However, we observed a monotonic increase in both decision criterion and task-evoked pupil dilation across the three paradigms, suggesting that the uncertainty of stimulus is not a primary factor responsible for adaptive decision making.

What behavioral state did our experimental conditions manipulate in the animals? For the paradigms with a lower probability of S+ trials, we observed a slowdown of reaction times and less impulsive licking. So it is possible that the animals were less engaged or less attentive in the task for these paradigms. Task engagement and attention have been show to modulate neural representation and cognitive processing in behavioral tasks (Eldar et al., 2013; Baruni et al., 2015; Luo and Maunsell, 2015; Lapborisuth et al., 2022), and thus might be an important contributor to adaptive decision making. However, the animals’ behavioral performance does not support this possibility because there was no significant difference in perceptual sensitivity across the three paradigms. But less engagement and poor attention usually lead to poor performance in perceptual tasks. Moreover, the task-evoked pupil dilation is largest in the paradigm where the probability of S+ trials was 20%, but previous work suggested larger task-evoked pupil dilation during more task-engaged or attentive periods (Hoeks and Levelt, 1993; Cardoso et al., 2019). Taken together, the behavioral adaptation during perceptual decision making in our experiments is unlikely to be primarily due to changes in behavioral states such as attention or engagement in the task. A possibility is that this behavioral adaptation is driven by different internal models involving probabilistic inference and expectation (Bouret and Sara, 2005; Yu and Dayan, 2005; Tervo et al., 2014). Future work with electrophysiological recordings in higher-order brain regions and neuromodulatory systems will help answer this intriguing question.

Our findings provide new evidence that adaptive decision making is dependent upon pupil-linked arousal. Previous work suggested that pupil size is able to reliably index the activation of the LC-NE system, as microstimulation of the LC evoked dramatic dilation of pupil in rats and non-human primates (Joshi et al., 2016; Liu et al., 2017) (but also see (Megemont et al., 2022)). Recent experimental results also demonstrated that pupil size co-varies with cholinergic activity in the brain to a lesser degree. Cholinergic neurons of the basal forebrain are more active during pupil dilation (Nelson and Mooney, 2016). 2-photon imaging revealed a positive correlation between the activation of cortical cholinergic axons and pupil size (Reimer et al., 2016). Recent results showed that phasic stimulation of serotonergic neurons in the dorsal raphe nucleus causes pupil size changes in mice performing a foraging task (Cazettes et al., 2021). Therefore, it is likely that the three neuromodulatory systems collectively exert influences on adaptive decision making in our experiments. It is intriguing for future studies to use selective manipulation to tease apart the contribution that each of the neuromodulatory systems provides to adaptive decision making.

## ACKNOWLEDGEMENTS

This work was supported by NIH R01MH112267, R01NS119813, and R21MH125107.

## AUTHORS CONTRIBUTION

Q.W. designed the study. S.N. and Q.W. performed the experiments. S.N. and B.J.S. analyzed the data. Q.W. wrote the manuscript. S.N and B.J.S. commented on the manuscript.

## DATA AVAILABILITY

All data and code are available upon request.

## DISCLAIMER

Q.W. is the co-founder of Sharper Sense, a company developing methods of enhancing sensory processing with neural interfaces.

